# The evolution of a condition-dependent mutation rate enhances evolvability

**DOI:** 10.64898/2026.07.09.737419

**Authors:** Timo J.B. van Eldijk, Jana M. Riederer, G. Sander van Doorn, Franz J. Weissing

**Affiliations:** Groningen Institute for Evolutionary Life Sciences, University of Groningen, Groningen, The Netherlands; Digital Competence Centre, Center for Information Technology, University of Groningen, Groningen, The Netherlands

**Author notes:** These authors contributed equally to the manuscript (shared first authorship) (Timo J.B. van Eldijk, Jana M. Riederer).

**Keywords:** plasticity, adaptive potential, individual-based simulation, theoretical model, adaptation to changing environments

## Abstract

Empirical studies have demonstrated that mutation rates may change with individual condition, such as in the case of stress-induced mutagenesis. This has led to the hypothesis that condition-dependent (or “plastic”) mutation rates could be selectively favoured, as the increased production of new mutants in times of maladaptation enhances evolvability, the ability to undergo adaptive evolution. However, while empirical evidence for condition-dependent mutation rates is accumulating, theoretical models studying their evolution are lacking. Here, we employ an individual-based simulation approach to examine the evolution of condition-dependent mutation rates in a changing environment. We find that condition-dependent mutation rates consistently evolve when the environment changes at an intermediate pace. Furthermore, populations with condition-dependent mutation rates are substantially better adapted to their (changing) environment. Finally, the evolutionary dynamics of condition-dependent mutation rates are both accelerated and destabilised when the mutation rate is self-referential (i.e., when mutator loci affect their own mutation rate). We conclude that condition-dependent mutation rates (and thus evolvability) can readily evolve in changing environments.

**Significance statement:** Mutation provides the raw material for evolution. Mutation rates thus tune evolvability, the ability to undergo adaptive evolution: if mutation rates are too low, evolution is impeded; if mutation rates are too high, adaptive traits cannot be maintained. Using a theoretical model, we explore the evolution of plastic mutation rates that systematically depend on the condition of the organism and its environment. An example is stress-induced mutagenesis in bacteria, which is implicated in the evolution of antibiotic resistance. We show that plastic mutation rates readily evolve, providing “well-timed” variation specifically when organisms are poorly adapted. Such plastic mutation rates thus facilitate better adaptation to changing environments, and their evolution provides an example of evolvability itself evolving.

## Introduction

The study of mutational processes is crucial to understanding evolution, as mutation provides the variation upon which selection acts. Since mutation rates affect what and how much genetic variation is available, they shape the ability of biological systems to undergo adaptive evolution. In other words, mutation rates are key determinants of evolvability (Bedau & Packard, 2003; Jones et al., 2007; Riederer et al., 2022).

The rate of mutation is subject to evolution. This is illustrated by the spread of mutator strains in the Lenski long-term evolution experiment, resulting in bacterial strains with elevated mutation rates (Sniegowski et al., 1999). Many researchers have studied the evolution of mutation rates both in models (Sniegowski et al., 2000; Bedau & Packard, 2003; Andre & Godelle, 2006; Desai et al., 2007; Lynch et al., 2016) and empirically, for example by using evolution experiments (Sniegowski et al., 1999; Colgrave & Collins, 2008; Couce et al., 2017; Sprouffske et al., 2018). In general, the optimal mutation rate strongly depends on the population’s “degree of adaptation”. When a population is close to a fitness peak, a low mutation rate is advantageous, as most mutations are deleterious. However, when a population is far from a fitness peak, an increased mutation rate can be beneficial, as it leads to the more rapid production of better-adapted variants. Alleles that increase the mutation rate can “hitchhike” along with any beneficial variants they produce. This is especially effective in asexual populations where the linkage between mutator alleles and the adaptive mutations they produce can be more easily maintained.

Some organisms are known to exhibit different mutation rates dependent on their condition. An example of this is stress-induced mutagenesis: various stressors are known to result in an increased mutation rate. There is widespread empirical evidence for such stress-induced mutagenesis in bacterial species, such as *Escherichia coli, Bacillus subtilis* and *Staphylococcus aureus* (Cirz et al., 2007; Foster, 2007; Debora et al. 2010; Ram & Hadany, 2012; Ha & Edwards, 2021). A stress-related elevation of the mutation rate has also been demonstrated in sexually reproducing eukaryotes, such as *Saccharomyces cerevisiae* and *Drosophila melanogaster* (Heidenreich, 2007; Agrawal & Wang, 2008) and even in human cancer cells (Cipponi et al., 2020). Stress-induced mutagenesis may be a non-adaptive side effect of stressful conditions (Ram & Hadany 2012). For instance, a stressor or stress response could impact DNA or the enzymes responsible for maintaining and replicating DNA, thereby leading to an increase in mutation rates, a phenomenon more accurately described as stress-associated mutagenesis (Bjedov et al., 2003; Tenallion et al., 2004; Mac Lean et al., 2013). There are, however, at least two adaptive explanations for stress-induced mutagenesis. First, when an organism is stressed, the costs of maintaining replicative fidelity could outweigh the benefits, inducing selection to switch off fidelity-enhancing mechanisms under stressful conditions (Ram & Hadany 2012). Second, stress may indicate that the organism is not well-adapted to its local conditions. As mentioned above, an elevated mutation rate may facilitate adaptation under such circumstances (Sniegowski et al., 1999; 2000).

We here explore the validity of this second explanation, which is based on the idea that selection can shape mutation rates in such a way that they fit the level of adaptation: in times of low stress (= a high level of adaptation), the organism benefits from a low mutation rate, whereas in times of high stress (= a low level of adaptation), it would benefit from an elevated mutation rate. According to this logic, it would be optimal to regulate the mutation rate in a condition-dependent manner: low under favourable conditions and high under unfavourable conditions. Here, “condition” refers to the state of the organism in relation to the current environment, which, by mechanisms such as stress-response systems, may indicate low levels of adaptation.

Whilst the empirical evidence for condition-dependent mutation rates is abundant, the condition dependence of mutation rates is much less studied from a theoretical point of view. The earliest models focused exclusively on the effect of deleterious mutations under condition-dependent mutation rates (Agrawal 2002; Baer 2008; Shaw & Baer 2011). In contrast, Ram and Hadany (2012, 2014, 2019) and Ram and colleagues (2018) also considered beneficial mutations and showed that stress-induced mutagenesis might be favoured by selection under a wide array of circumstances. However, the models by Ram and Hadany (2012, 2014, 2019) assume that the loci determining the relationship between an individual’s condition and its mutation rate are fixed and not evolvable. In other words, they do not address the evolutionary emergence of the relationship between an individual’s condition and its mutation rate.

Here, we use an individual-based simulation approach to better understand the evolution of condition-dependent mutation rates. We consider a population that is constantly adapting to a changing environment. This adaptation is mediated by a phenotypic trait that is subject to both beneficial and deleterious mutations. In contrast to the studies of Ram and Hadany (2012, 2014, 2019), we focus on the evolutionary dynamics of the relationship between individual condition and the mutation rate. To this end, we treat the mutation rate as a plastic trait that responds to an individual’s condition according to a threshold-shaped reaction norm.

We address two questions. First, can condition-dependent mutation rates evolve, and, if so, what circumstances are favourable for their evolution? In particular, how is the evolution of condition-dependent mutation rates related to environmental change? Second, how do condition-dependent mutation rates affect the speed of adaptation in a changing environment? In other words, to what extent does a condition-dependent mutation rate enhance the evolvability of a population?

Most previous models of the evolution of mutation rates assume (often implicitly) that the mutator loci (i.e., the loci controlling the mutation rate) mutate according to an externally given fixed mutation rate (e.g., Agrawal 2002, Baer 2008; Shaw & Baer 2011; Ram et al., 2018; Ram & Hadany 2012, 2014, 2019). Here, we also consider the option that the mutation rate at the mutator loci is “self-referential” in the sense that this mutation rate is affected by the mutator loci themselves.

## Methods

### Model overview

We use an individual-based simulation approach to study the evolution of a population in an environment that may randomly change from one generation to the next. Each individual has a genetically determined phenotype *P*. The expected number of offspring of an individual is positively related to the degree to which the individual’s phenotype *P* matches the current state of the environment *E*. Figure 1A illustrates the change in the environment and the tracking of the environment by the average phenotype in the population due to the joint action of mutation, selection, and genetic drift.

**Figure 1.**
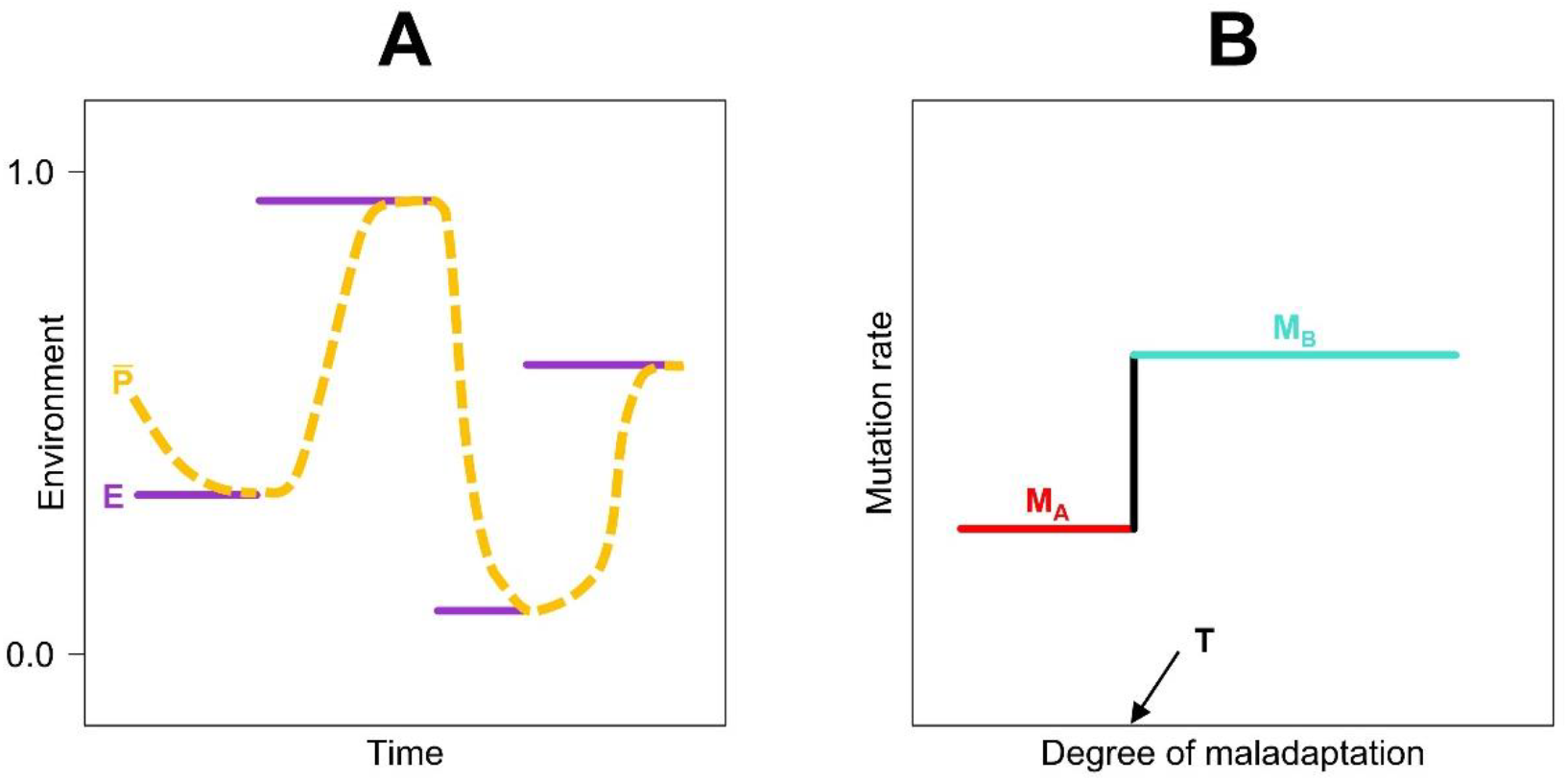
Schematic illustration of the model setup. **(A)** We consider a randomly changing environment E (purple line segments) that switches to a new state once in a while. The probability that such a switch occurs is constant and low, so that the environment tends to remain stable for a while. During such periods, natural selection acts to reduce the mismatch between the genetically determined phenotype of individuals and the current environment. As a result, the mean phenotype P tends to track the environment. **(B)** Our model allows mutation rates to be condition-dependent. In this case, the mutation rate of an individual is governed by three heritable traits (M_A_, M_B_, T): the mutation rate M_A_ is expressed if the degree of maladaptation is smaller than the threshold T; otherwise, M_B_ is expressed. The “degree of maladaptation” is quantified by 1 – F(P,E), the difference between maximal fitness and the fitness F(P,E) of phenotype P in the current environment E, and it is positively related to the mismatch |P–E| between P and E.

We consider two scenarios of mutation rate evolution: the evolution of a constitutive mutation rate and the evolution of a condition-dependent mutation rate. In the first scenario, the mutation rate is determined by the allele at a mutator locus; this allele is also inherited from parent to offspring, subject to mutation. In the second scenario, the mutation rate is dependent on the condition of the individual, where the individual’s “condition” is quantified by its degree of (mal)adaptation, i.e., it reflects how well the individual’s phenotype *P* matches the current state of the environment *E*. As illustrated in Figure 1B, condition-dependent mutation rates are determined by a reaction norm encoded by three loci: the first locus codes for the threshold value of the reaction norm – this will set the individual’s bench mark for “deciding” whether it is in a “bad” or in a “good” condition; the other two loci determine, respectively, the mutation rates expressed on either side of this threshold. That is: the allele on mutator locus *A* determines the mutation rate when in good condition, while mutator locus *B* determines the mutation rate in bad condition. Again, the alleles at all three loci encoding the reaction norm are transmitted from parent to offspring, subject to mutation.

In both scenarios, the alleles at the mutation-relevant loci are also affected by mutation. We consider two variants for the mutation rate at these loci: (a) a constant, externally given (and hence not evolving) mutation rate or (b) a “self-referential” mutation rate, which is identical to the mutation rate at the locus determining the phenotype *P* (and hence determined by the mutation-relevant loci themselves). Note that throughout, we model the evolution of mutation *probabilities*, i.e., the probability of a mutation occurring during a reproduction event. However, in keeping with conventional terminology, we refer to these mutation probabilities as “mutation *rates*”.

### Model assumptions

#### Environmental change

The environment is characterised by a real number *E*, ranging from 0 to 1. The environmental state *E* remains constant throughout a generation and is the same for all members of the population. From one generation to the next, *E* changes with probability χ. Hence, the expected duration of environmental stasis is *D* = 1/χ. Whenever environmental change occurs, a new environmental state is drawn from the uniform distribution on the interval [0,1]. Unless stated otherwise, we chose χ = 0.01, or equivalently *D* = 100 generations.

#### Phenotypes, mismatch, and fitness

Each individual has a genetically determined phenotype *P* (see below), represented by a number between 0 and 1. The expected reproductive success (“fitness”) *F(P,E)* of individuals with phenotype *P* in an environment in state *E* is negatively related to the mismatch |*P–E*| and proportional to the Gaussian

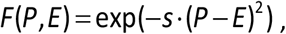

where the parameter *s* measures the strength of selection. For all simulations reported in the main text, we chose *s* = 10, corresponding to a standard deviation of 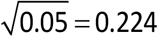 of the Gaussian (but see the Supplement for weaker selection, i.e. lower values of *s*). Note that since P and E are elements of the unit interval [0,1], so are the mismatch |*P–E*| and *F(P,E)*.

#### Selection and reproduction

We consider a population with discrete, non-overlapping generations and a fixed population size of *N* individuals. In all simulations reported, we chose *N* = 1,000. Inheritance is asexual; hence, offspring inherit their parent’s phenotype unless a mutation occurs. A new generation is produced as follows: for each of the *N* offspring individuals, a parent is drawn from the previous generation by means of a weighted lottery (with replacement), where the probability of an individual *i* with phenotype *P*_*i*_ to be chosen is proportional to the individual’s fitness *F*(*P*_*i*_,*E*). This procedure ensures that the expected number of offspring of each individual is proportional to *F*(*P*_*i*_,*E*).

#### Inheritance

Individuals are haploid and have either two or four gene loci (depending on the simulation scenario), each harbouring infinitely many alleles (corresponding to real numbers). All parental alleles are transmitted from parent to offspring, subject to mutation. One locus encodes the phenotype *P* of an individual, while the other loci determine the mutation rate at the *P* locus. The alleles at the *P* locus are elements of the unit interval [0,1]; each allele directly corresponds to the phenotype it encodes. In the case of a constitutive mutation rate, the second locus encodes the mutation rate *M* at the *P* locus. To be more specific, the alleles at the *M* locus are elements of the unit interval [0,1]; each allele directly specifies the mutation rate. In the case of a condition-dependent mutation rate, the mutation rate at the *P* locus is determined by three parameters (*M*_*A*_,*M*_*B*_,*T*), as indicated in Figure 1B. *M*_*A*_ and *M*_*B*_ are again elements of the unit interval [0,1]. The alleles at the threshold locus *T* are limited to the interval [−0.1,1.1]. The reason for this is as follows: if 0<*T*<1, both *M*_*A*_ and *M*_*B*_ may be expressed, depending on the individual’s condition. To ensure that evolutionary outcomes where only *M*_*A*_ or only *M*_*B*_ is expressed are possible, we limit *T* not to the unit interval [0,1] but to the larger interval [−0.1,1.1]. This allows us to distinguish the case of a low but positive threshold, which indicates a condition-dependent mutation rate in which individuals switch between *M*_*A*_ when in good condition vs. *M*_*B*_ when in suboptimal condition, from the case of a low and negative threshold, which indicates that *M*_*B*_ is expressed constitutively (as the degree of maladaptation can never be negative).

#### Mutation

Whenever a new offspring is produced, it inherits the parental genome, subject to mutation. If a mutation occurs, the new trait value is given by *Value*_*new*_ *= Value*_*old*_ *+ δValue*, where the mutational step size *δValue* is drawn from a normal distribution with mean zero and standard deviation 0.05. Values at the *P* and *M* loci are limited to [0,1]: thus, if a value of *δP* or *δM* is such that *P*_*new*_ or *M*_*new*_ is outside of [0,1], *δP* or *δM* is redrawn. Values at the T locus are limited to [−0.1,1.1]: thus, if a value of *δT* is such that *T*_*new*_ is outside of [−0.1,1.1], *δT* is redrawn. At the *P* locus, a mutation occurs with a probability that is determined by the mutation-related loci. In the model variant with externally given mutation rates for mutator loci, the *M* and *T* loci (which encode the mutation rate) mutate with a constant probability *μ* = 0.001. In the model variant with self-referential mutation rates, the mutation rates at the *M* and *T* loci are equal to that at the *P* locus (i.e., the mutation-rate-determining loci determine their own mutation rates).

#### Initialisation

All simulations start with a monomorphic population, with all individuals having allele *P* = 0.5 at the locus determining the phenotype. Unless stated otherwise, all mutation rates are initialised at the value 0.001, and the threshold locus is initialised at *T* = 0.5.

#### Simulation details

All simulations were run for 200,000 generations, as most replicates reached stable values of the mutation rates (aside from noise inherent to a system with evolving mutation rates) within this timeframe. For each initialisation and parameter setting, 100 replicate simulations were run. The simulation programme was written in C^++^, and the simulation data were processed and analysed in *R*.

#### Model variants

An earlier version of this study is documented in Chapter 4 (p. 89-122) of the PhD thesis of T.J.B. van Eldijk (2024). This earlier version is similar to the present one in most respects, except that the mutation process is implemented differently. Reassuringly, this different implementation leads to broadly similar conclusions.

## Results

### Evolution of constitutive mutation rates

Figure 2 illustrates the evolution of constitutive (i.e., non-plastic) and non-self-referential mutation rates. Figure 2A shows how the evolutionary outcome depends on the average time of environmental stasis *D*. The mutation rate (or more precisely, the mutation probability per reproduction event) evolves to high values in rapidly changing environments (such as *D* = 10; or equivalently, rate of environmental change χ = 0.1;) and approaches zero in slowly changing or constant environments (*D* ≥1000; χ ≤0.001). The increase in mutation rate with the rate of environmental change is not unexpected, as a high mutation rate allows evolution to keep pace with rapid environmental change. Figure 2B illustrates, for our default value *D* = 100, the evolutionary trajectories and the convergence of different replicates to the same equilibrium value, with considerable fluctuations around the equilibrium. To verify that a higher mutation rate indeed leads to better adaptive tracking of a rapidly changing environment, we investigated the average mismatch between phenotype and environment across a range of fixed (i.e., non-evolving) constitutive mutation rates, for the case *D* = 100 (Figure 2C). The mismatch between phenotype and environment is highest for very low and very high mutation rates and minimal at a mutation rate of about 0.1, with a steep increase in average mismatch when the mutation rate decreases, but only a shallow increase in mismatch when the mutation rate increases from 0.1. This is consistent with Figure 2A, which shows that for *D* = 100, mutation rates evolved to be close to 0.1.

**Figure 2.**
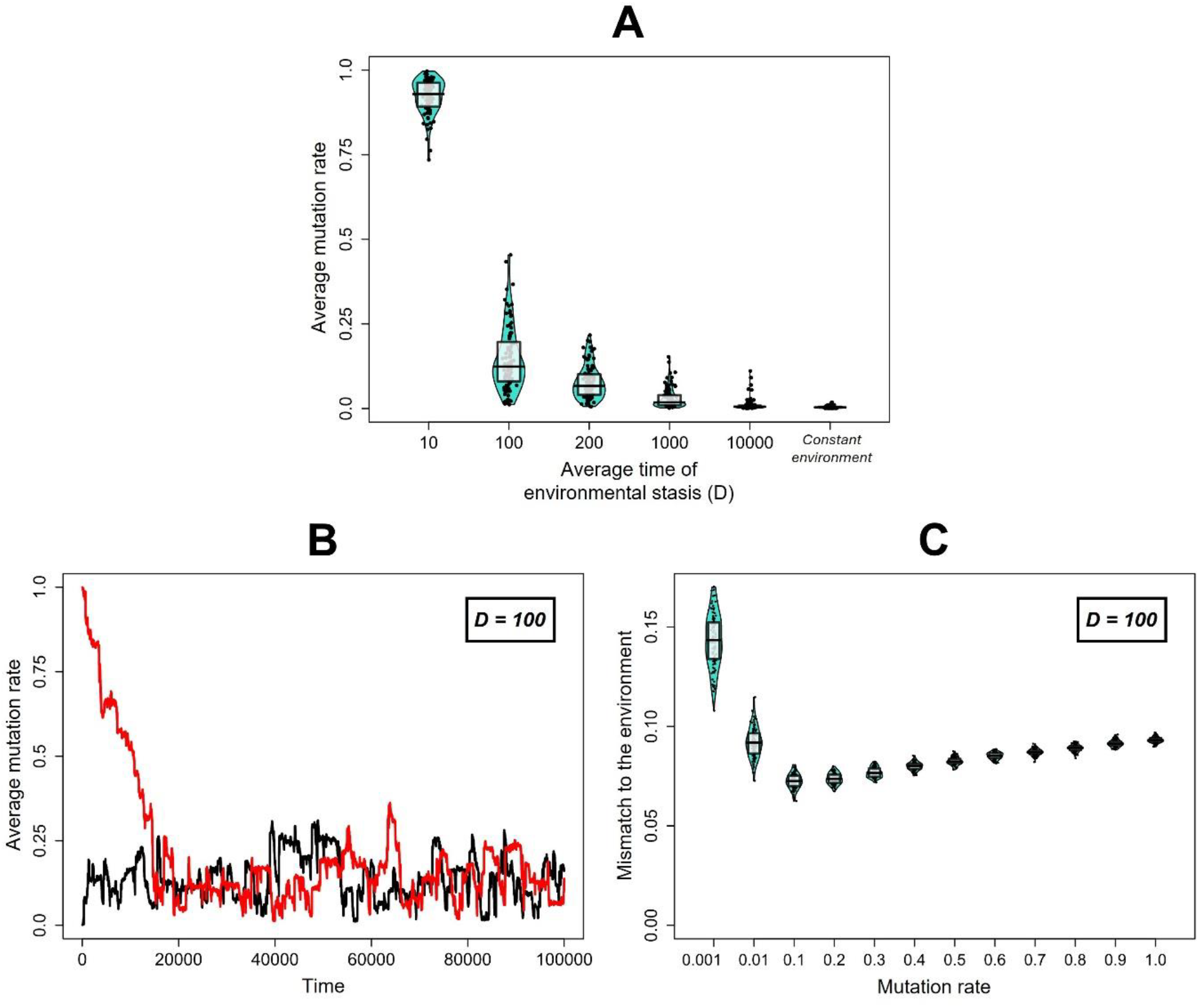
Evolution towards an optimal constitutive mutation rate, and its dependence on the rate of environmental change. Panel **(A)** shows the mutation rate that evolved under different rates of environmental change for six values of D=1/χ. Each violin summarises 100 replicate simulations. Here, each point corresponds to the average mutation rate evolved at the end of a replicate simulation (i.e., in the 200,000th generation). In panel **(B)**, a representative replicate (in black) shows a typical evolutionary trajectory of the population-average mutation rate, evolving in an environment that changes at an intermediate rate (D = 100) for 100,000 generations. For comparison, a second replicate (initialised with a mutation rate of 1.0 rather than the default of 0.001) is shown in red; the two replicates converge to similar average mutation rates. Panel **(C)** shows the average mismatch between phenotype and an environment that changes at an intermediate rate (D = 100), for different values of a non-evolving constitutive mutation rate. Each point represents the time-average mismatch calculated across all individuals over the last 10,000 generations of a simulation (out of 200,000 generations). Each violin summarises the results of 100 replicate simulations, with the solid lines indicating the median and the boxes indicating the interquartile range.

The outcome of the evolution of a constitutive self-referential mutation rate is similar to that of a non-self-referential one (Figure 3B): across replicates, higher mutation rates evolve under faster environmental change. However, individual evolutionary trajectories exhibit rapid fluctuations (Figure 3A), and there is greater variation between replicates (Figure 3B) than with a non-self-referential mutation rate.

**Figure 3.**
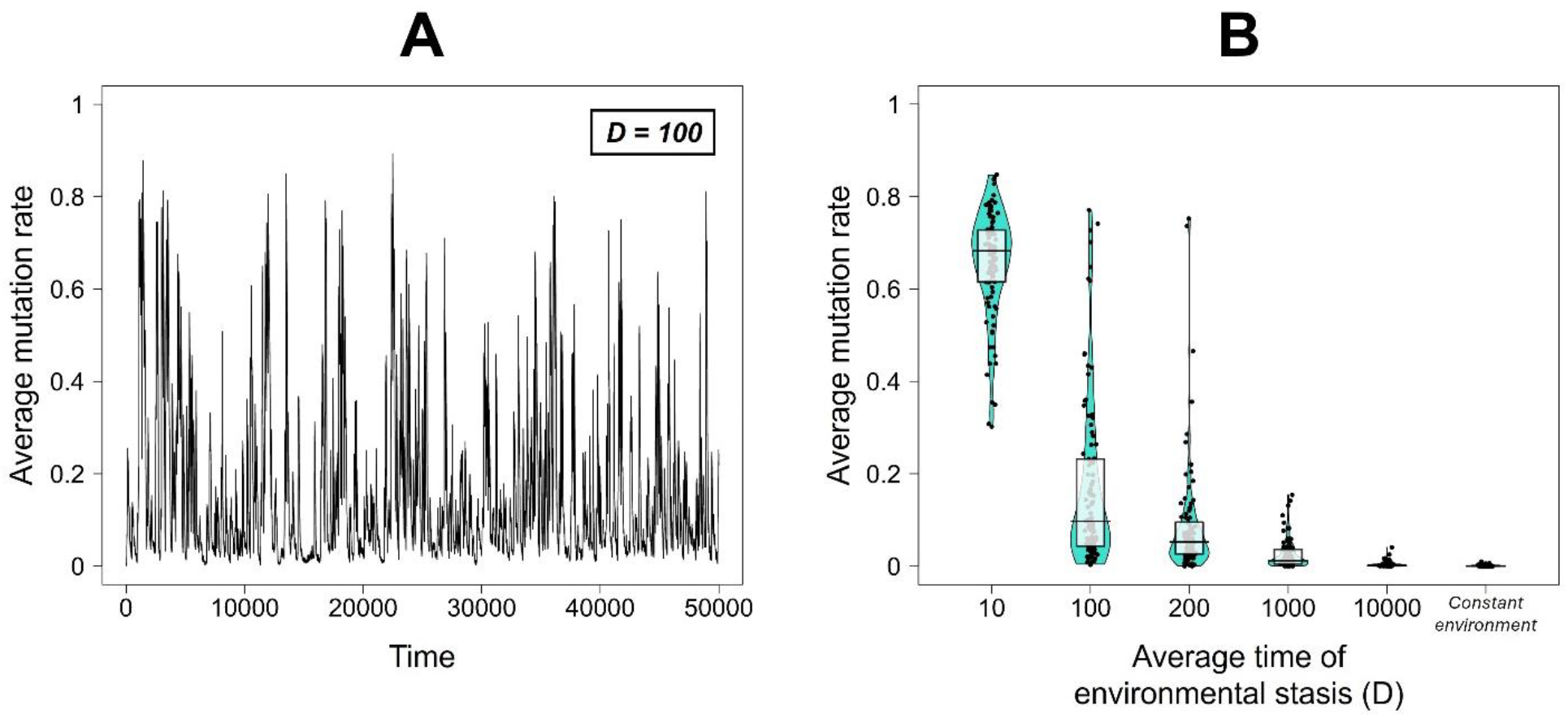
Evolution of a constitutive mutation rate under different rates of environmental change when the mutation rate is self-referential. This figure is similar to Figure 2 panels A and B, but displays the results for a self-referential mutation rate. Panel **(A)** shows the population-average mutation rate of a single representative replicate over 50,000 generations (again for the default case of D = 100), which exhibits rapid fluctuations. Panel **(B)** shows the mutation rates that evolved under different rates of environmental change. For six values of D=1/χ, a violin plot summarises the results of 100 replicate simulations after 200,000 generations, as in Figure 2B.

### Evolution of condition-dependent mutation rates

For our default setting *D* =100, Figure 4 shows the outcomes of 100 replicate simulations that allowed the evolution of condition-dependent mutation rates. All simulations were initialised with mutation rates *M*_*A*_=*M*_*B*_=0.001 and a threshold value *T=0*.*5*. Over the course of evolution, *M*_*A*_ evolved to low values (Figure 4A), whereas *M*_*B*_ evolved to high values (Figure 4B). The threshold *T* evolved to be low but positive (Figure 4C). For a representative replicate, see Figure 4D. Together, these values form a reaction norm (Figure 4E): in good condition (i.e., when the degree of maladaptation is low), the low mutation rate *M*_*A*_ is used, but once condition worsens (i.e., when the degree of maladaptation increases), the high mutation rate *M*_*B*_ is used. The mutation strategy (*M*_*A*_, *M*_*B*_, *T*) that had evolved at the end of the simulation is shown in Figure 4F: all replicates evolved towards a low *M*_*A*_ and a high *M*_*B*_. Whether this combination corresponds to a plastic, condition-dependent mutation rate depends on the (average) threshold value 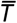 and is indicated by the colour code: when *T* is between zero and one, individuals can switch between mutation rates depending on their condition. Out of 100 replicates, 99 evolved condition-dependent mutation rates (0<T<1, colour blue – yellow), and one evolved to constitutively express mutation rate *M*_*B*_ (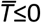, colour black). Of the 99 replicates showing condition-dependency, 98 evolved low values of 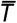: here, mutation rates plastically switch from low to high values for even a small decrease in individual condition.

**Figure 4.**
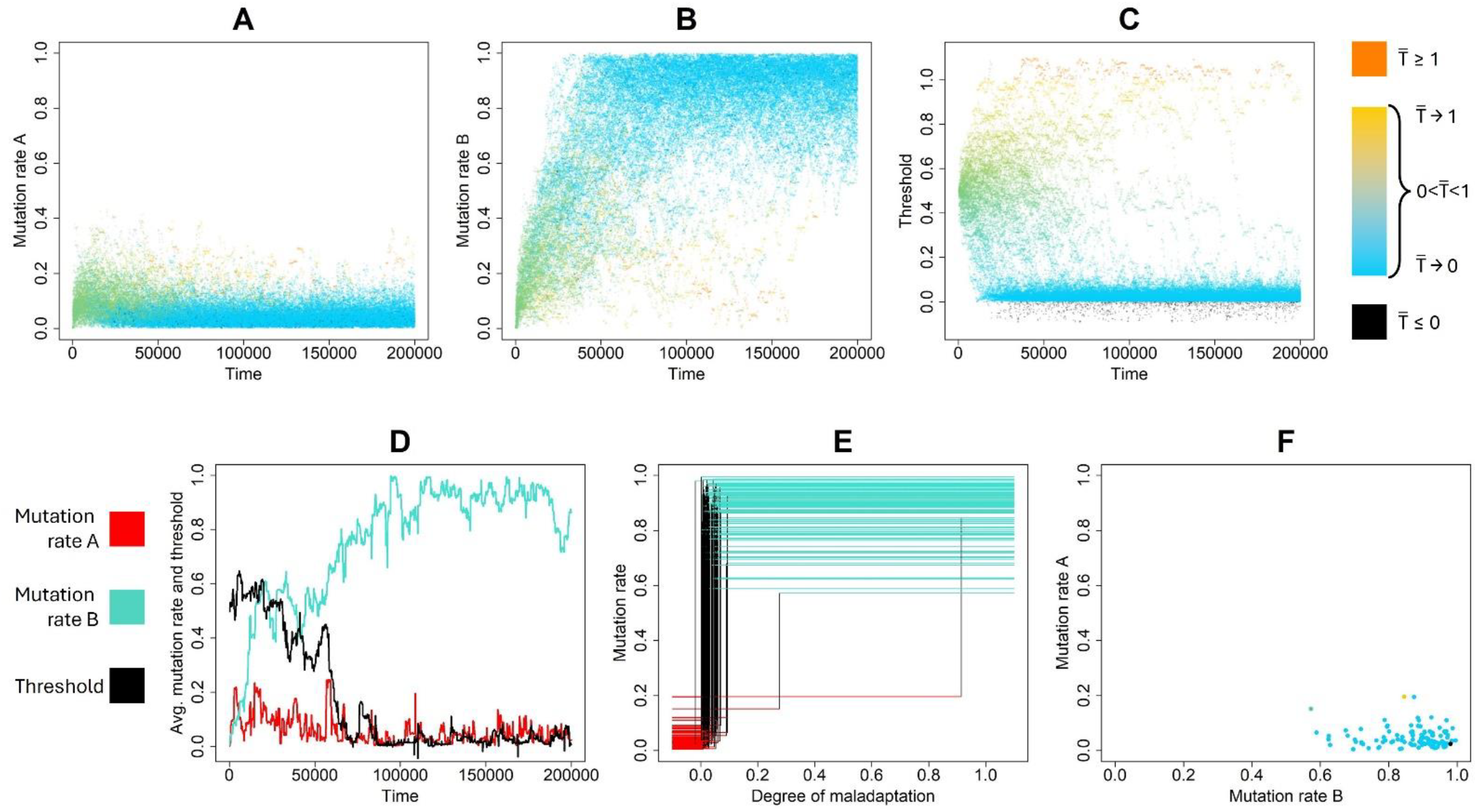
Evolution of condition-dependent mutation rates. Panels **(A), (B)** and **(C)** show time series of the evolution of the three mutator loci, for 100 replicate populations. Points show the population average at a particular timepoint for a particular replicate, and are colour-coded based on their corresponding population average 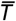: values evolving for low but positive threshold values appear blue, while rates evolving for threshold values that are high but below one appear yellow; rates evolving for 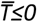 are coloured black and rates evolving for 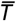≥1 are orange. Starting from the same initial values of M_A_=M_B_=0.001, the mutation rate M_A_ evolved to be low (A) whilst the mutation rate M_B_ evolved to be high (B). Starting from the initial value of 0.5, the threshold T evolved to be low but positive in 98 out of 100 replicates (C). Panel **(D)** shows these dynamics for a representative simulation. Here, the population averages of M_A_, M_B_, and T are shown in red, turquoise, and black, respectively. Panel **(E)** shows, for the same 100 replicates as shown in (A)-(C), the population average reaction norm after 200,000 generations, i.e., at the end of the time series shown in (A)-(C). The colour code is as in (D). **(F)** shows which values of mutation rate M_A_ co-evolved with which values of mutation rate M_B_, again for the same 100 replicate populations. Each point indicates a pair of average mutation rate values evolved at the end of one replicate simulation (i.e., in the 200,000th generation); colour code as in (A) – (C). For all results shown in this figure, the average duration of environmental stasis is D = 100.

In the Supplement, we present corresponding simulations, but for weaker selection (i.e., lower values of *s*). For an intermediate selection strength (*s*=1, Figure S1), a condition-dependent mutation strategy (*M*_*A*_, *M*_*B*_, *T*) still evolves, albeit more slowly and in a more stochastic fashion. This is no longer the case when selection is very weak (*s*=0.1, Figure S2) and apparently dominated by genetic drift.

### Effect of the environmental change rate on condition-dependent mutation rates

Figure 4 considered the default setting *D* = 100, where the environment remains constant for, on average, 100 generations. Figure 5 shows the outcome of 100 replicate simulations for six different values of *D*. Let us start with the default case of *D* = 100. Similar to what we saw in Figure 4, condition-dependent mutation rates evolve in all 100 replicates: threshold values *T* are generally low but positive (Figure 5C), such that *M*_*A*_ is expressed when in good condition, and *M*_*B*_ is expressed otherwise. *M*_*A*_ consequently evolves to be low (Figure 5A), whereas *M*_*B*_ evolves to be high (Figure 5B). The same pattern can be seen as a single cluster of points in Figure 5E, as all replicates follow the same mutation strategy. *D* = 200 and *D =* 1000 exhibit a similar pattern, as condition-dependent mutation rates consistently emerge. Next, let us consider a fast-changing environment with *D* = 10. Here, the average thresholds evolved in the 100 replicate simulations are bimodally distributed, taking on either high values (including values above one) or low values (including values below zero, Figure 5C). Both *M*_*A*_ and *M*_*B*_ show a distribution biased towards high values but spanning their entire possible range (Figure 5A,B). Figure 5D shows that this translates into two different “mutation strategy” clusters. The first cluster corresponds to the cloud of points that are coloured yellow or red (indicating that 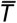 is close to or above one). The corresponding populations evolved to constitutively express *M*_*A*_: they are characterised by high values of *M*_*A*_, and high values of *T*; here, *M*_*B*_ is seldom, if ever, expressed, and thus mainly evolves via genetic drift. The second cluster corresponds to the cloud of points coloured in blue or black (indicating that 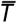 is close to or below zero). These populations evolved to largely express *M*_*B*_: they are characterised by high values of *M*_*B*_, and low values of *T*; here, *M*_*A*_ is seldom expressed, and thus drifts. Thus, under rapid environmental change, condition-dependent mutation rates do not consistently emerge. Instead, when the environment changes rapidly, constant adaptation is necessary, selecting for a constitutive expression of a high mutation rate, which may be mutation rate *M*_*A*_ (cluster one) or mutation rate *M*_*B*_ (cluster two). Finally, when the environment remains constant or changes very rarely (every 10,000 generations), there is selection for a low mutation rate, leading to low values of *M*_*A*_ (see Figure 2). In this scenario, the mismatch between phenotype and environment will typically be very small, so that selection on *M*_*B*_ and *T* is very weak (as long as *T* is positive). Accordingly, *M*_*B*_ is expressed only rarely or not at all, and *M*_*B*_ and *T* will largely evolve by genetic drift (note that *T* remains limited to positive numbers though, Figure 5F).

**Figure 5.**
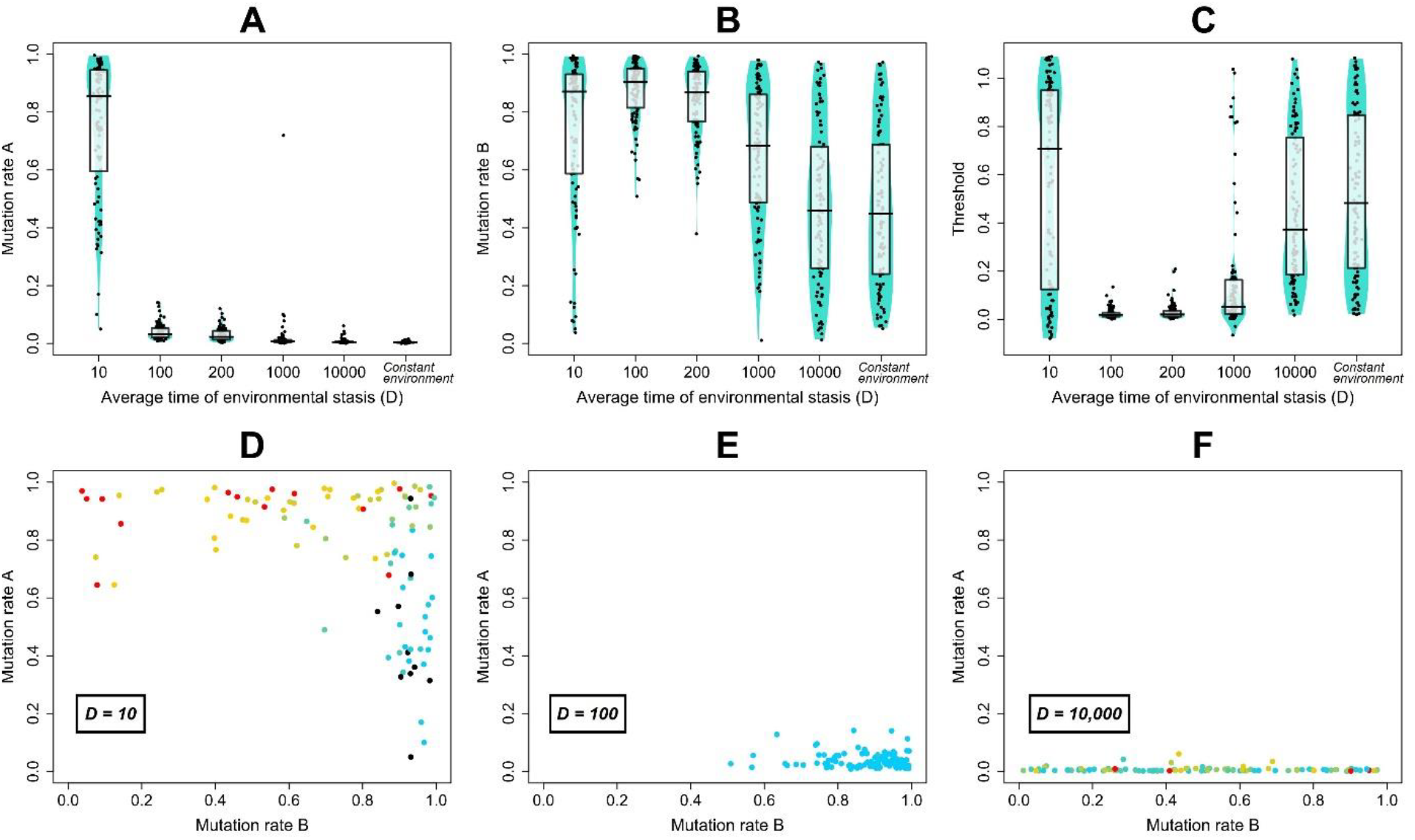
Effect of the average time of environmental stasis on the evolution of condition-dependent mutation rates. For six values of D=1/χ, three violin plots show **(A)** the population average of M_A_, **(B)** the population average of M_B_, and **(C)** the population average of T. Each violin shows the population averages of 100 replicate populations after 200,000 generations. All simulations were initialised at M_A_=M_B_=0.001 and T=0.5. Each point corresponds to one replicate population; the solid lines indicate the median. Panels **(D), (E)** and **(F)** show which values of M_A_ co-evolved with which values of M_B_, for D=10, D=100, and D=10,000, respectively, colour-coded based on 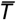, as in Figure 4.

### Self-referential mutation rates

In Figures 4 and 5, we assumed that the three loci determining the mutation rate mutate with a fixed mutation probability (μ = 0.001). Now, we consider a self-referential mutation rate: the loci encoding the mutation rate mutate with the same probability as the *P* locus (whose mutation rate is encoded by these loci). The resulting evolutionary dynamics are similar in several ways: again, *M*_*A*_ evolves to be low, *M*_*B*_ to be high, and *T* to be low but positive (Figure 6B, C). These values again form a clear reaction norm (Figure 6B), demonstrating the evolution of condition-dependent mutation rates: individuals in good condition (i.e., with a low degree of maladaptation) use the low mutation rate *M*_*A*_, but switch to using the high mutation rate *M*_*B*_ when their condition worsens (i.e., when the degree of maladaptation increases). However, the comparison of Figure 6 (self-referential mutation rate) with Figure 4 (fixed mutation rate at the mutator loci) also highlights some notable differences: for instance, in the case of self-referential mutation rates, there is more variation between the replicates (Figure 6B, C) and the evolutionary dynamics, though faster to establish, is less stable (Figure 6A). This matches the results for constitutive mutation rates, shown in Figures 2 and 3: self-referential mutation rates exhibit similar but “noisier” dynamics, both in the case of constitutive mutation rates and in the case of condition-dependent mutation rates.

**Figure 6.**
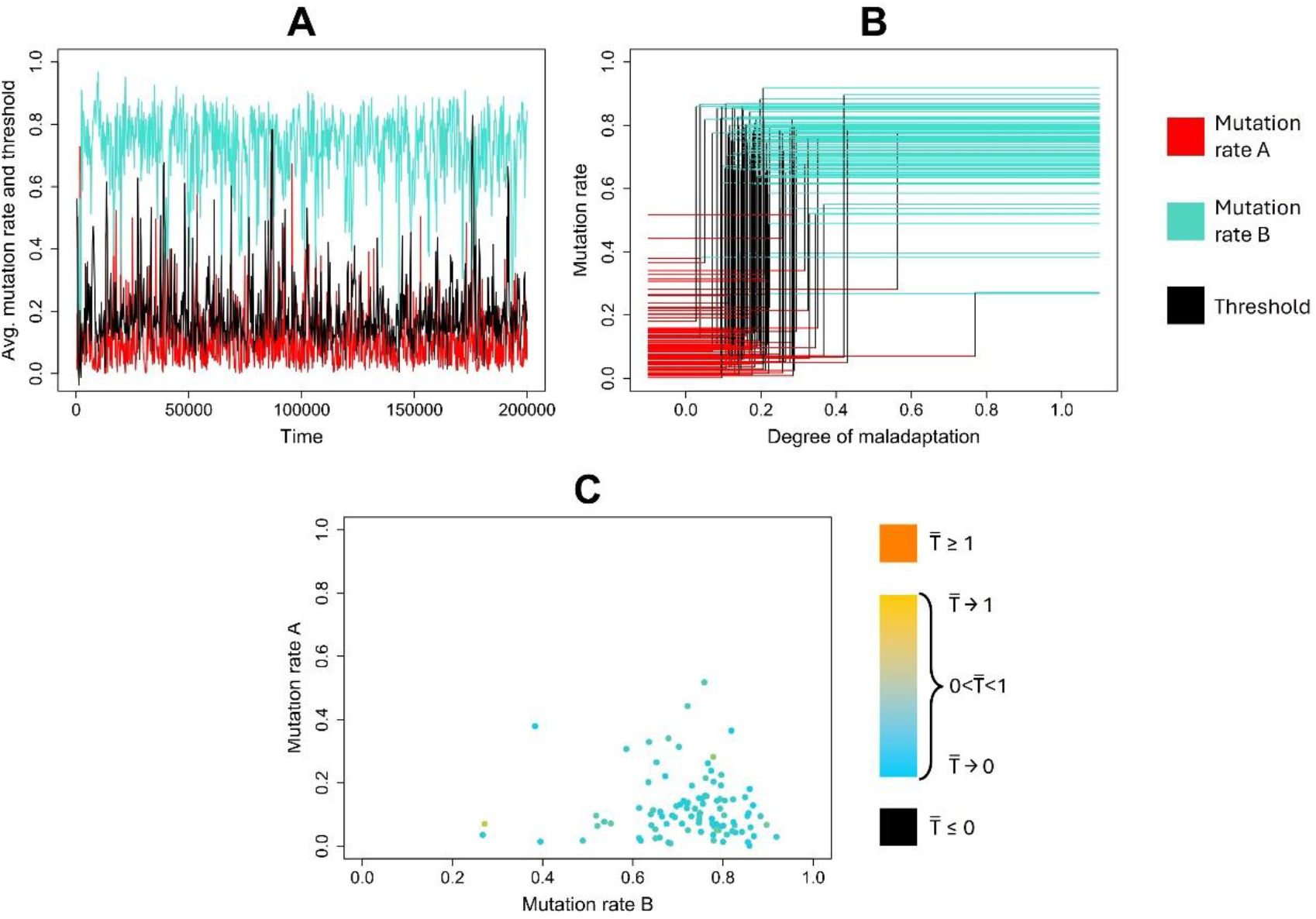
Evolution of condition-dependent mutation rates when the mutation rate at the mutator loci is self-referential. Panel **(A)** shows the evolutionary trajectories of the three mutator loci for a representative replicate population, colour-coded as in Figure 4D. Panel **(B)** shows, for 100 replicate populations, the population-average reaction norm after 200,000 generations. Colour code as in (A). Panel **(C)** shows which values of mutation rate M_A_ co-evolved with which values of mutation rate M_B_, again for the same 100 replicate populations, and colour-coded based on 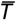, as in Figure 4. For all results shown in this figure, average duration of environmental stasis D = 100.

### Implications for adaptation in a changing environment

Different mutation strategies influence the adaptive tracking of a changing environment. Figure 7 shows how the time-averaged mismatch between phenotype and environment depends on the evolved mutation-rate strategy and on the speed of environmental change: when the environment changes at an intermediate speed (e.g. *D* = 100), condition-dependent mutation rates arise, which allows for better tracking of the environment (and consequently, a lower average mismatch to the environment, Figure 7A). However, when environmental change is very fast (*D* = 10) or very slow (e.g. *D* = 10,000), condition-dependent mutation rates do not consistently emerge (Figure 5D and Figure 5F), and the degree of mismatch to the environment does not differ from that observed in populations with constitutive mutation rates (Figure 7A). For the default setting of *D* = 100, we show the population average mismatch to the environment, for all four mutation strategies explored in this study: the constitutively expressed non-self-referential mutation rate, the constitutively expressed self-referential mutation rate, the plastic (i.e., condition-dependent) non-self-referential mutation rate, and the plastic self-referential mutation rate. Condition-dependent mutation rates show better adaptive tracking than constitutive mutation rates, both when self-and non-self-referential. Furthermore, a self-referential mutation rate leads to better adaptive tracking when it is constitutive, but not when it is condition-dependent.

**Figure 7.**
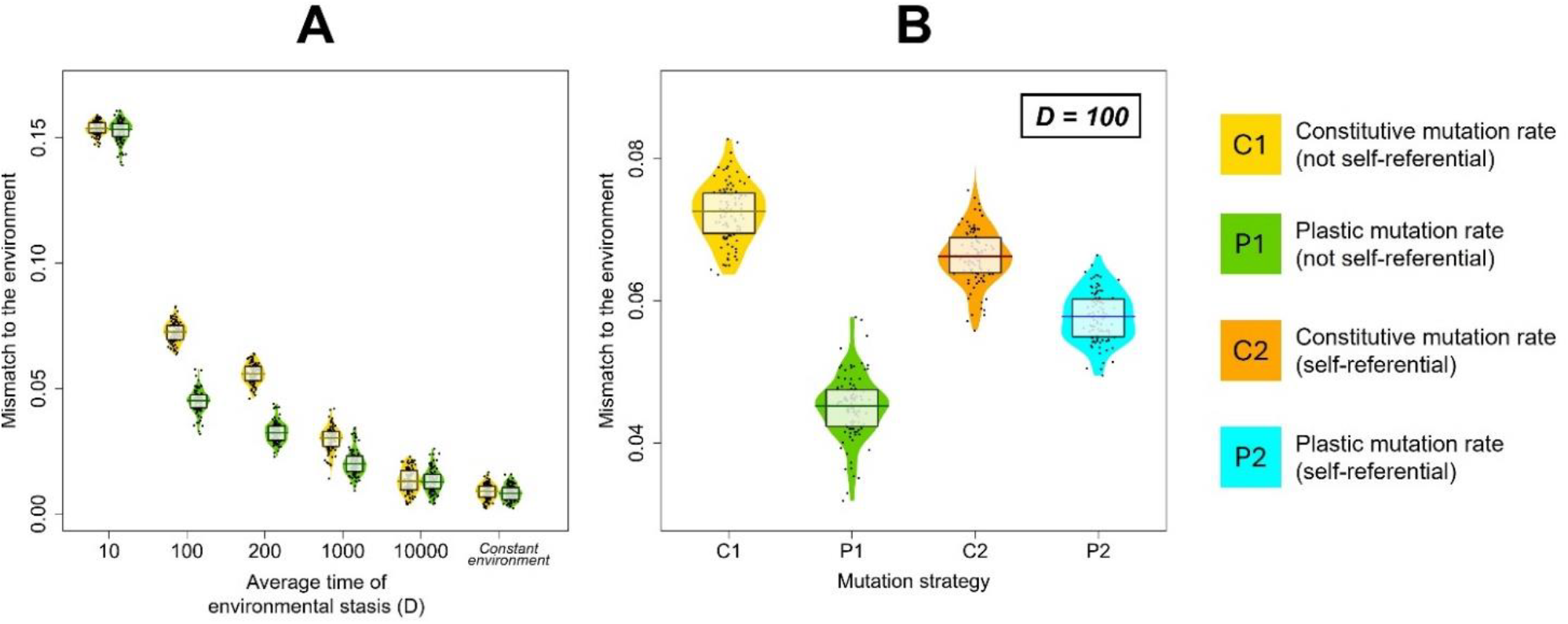
Time-averaged mismatch between phenotype and environment for different evolved mutation-rate strategies. Panel **(A)** shows the mismatch between phenotype and environment under different speeds of environmental change. Two mutation strategies are shown: a constitutive mutation rate (C1, yellow) and a plastic (i.e., condition-dependent) mutation rate (P1, green), both non-self-referential. For our default scenario of D = 100, panel **(B)** shows the mismatch between phenotype and environment for all four mutation strategies: the pair of violins on the left shows the results in the case of non-self-referential mutation rates, either when mutation rates are assumed to be constitutive (C1, yellow) or when mutation rates are allowed to evolve to be condition-dependent (i.e., plastic, P1, green). The pair of violins on the right shows the results for self-referential mutation rates, again either when assumed to be constituent (C2, orange) or when allowed to evolve to be condition-dependent (P2, blue). In both panels, each violin summarises the outcome of 100 replicate simulations; each point represents the time-average of the mismatch between phenotype P and environmental state E, calculated over all individuals in the last 10,000 generations of a simulation (out of 200,000 generations). In panel (B), two replicates of C2 lie outside the plotted range, as they exhibited a mismatch of 0.12 and 0.10, respectively.

## Discussion

Mutation rates are linked to a key concept in evolutionary biology: evolvability, the capability of a biological system to undergo adaptive evolution. Since mutation rates affect the rate at which variation is generated, they are a crucial determinant of evolvability (Badeau & Packard, 2003; Jones et al., 2007; Riederer et al., 2022). Thus, our results shed light on evolvability and its evolution. As a standard of comparison, we first considered the evolution of a constitutive mutation rate. Here, we found (in line with previous empirical studies, e.g. Sprouffske et al. 2018) that moderately elevated mutation rates facilitate the closest tracking of the changing environment – in other words, moderately high mutation rates are associated with the highest evolvability. The idea that an elevation in mutation rate can facilitate adaptation but only up to a point is well established (e.g., Baedau & Packard, 2003; Sprouffske et al. 2018; Matic et al., 2019): when mutation rates are low, there may be insufficient influx of variation; if mutation rates are high, newly gained adaptations may not be maintained. Optimal mutation rates thus face a trade-off between “exploration” or “flexibility” (the ability to find new adaptive variants) and “exploitation” or “robustness” (the ability to retain and thus make use of these variants in the face of mutation pressure; see Baedau & Packard, 2003; but see Andre & Godelle, 2006). Moreover, higher mutation rates evolved in environments that changed more rapidly: under rapid change, there is an increased need for phenotypic change (exploration/flexibility), whereas in stable environments, there is an increased need for phenotypic stability (exploitation/ robustness). Our findings are again in line with previous studies on mutation rate evolution (e.g. Ishii et al., 1989; Bedau & Packard, 2003; Tanaka et al., 2003; Palmer & Lipsitch, 2006; Wielgoss, 2013; Carja et al., 2014).

Next, we consider the evolution of condition-dependent mutation rates. Condition-dependency arose specifically in environments changing at an intermediate rate; in contrast, under rapid environmental change, constitutive high mutation rates evolved, and under fast environmental change, constitutive low mutation rates evolved. This can be viewed in light of the exploration-exploitation trade-off mentioned above: under rapid environmental change, selection for exploration predominates (selecting for high mutation rates); under slow environmental change, selection for exploitation predominates (selecting for low mutation rates); and under environmental change at an intermediate pace, the selection pressure alternates (selecting for condition dependency). In such an environment, a moderately elevated constitutive mutation rate and a condition-dependent mutation rate can thus be viewed as two solutions to the same problem: a moderately elevated constitutive mutation rate presents a compromise between the need for exploration and exploitation; condition-dependent mutation rates circumvent this issue by plastically switching between exploration (high mutation rates) and exploitation (low mutation rates) as needed. In line with this, we also find that condition-dependent mutation rates allow a closer tracking of the environment (again, under intermediate speeds of environmental change): when the environment changes fast enough that new adaptation is frequently needed, but slow enough to still necessitate phenotypic robustness, then condition-dependent mutation rates can increase evolvability. This echoes previous findings that stress-induced mutagenesis can enable crossing of fitness valleys, albeit via a mechanism different from the one we considered (Ram & Hadany, 2014).

Most models on the evolution of mutation rates implicitly assume that alleles at mutator loci mutate at a fixed, externally given rate (Agrawal 2002; Baer 2008; Shaw & Baer 2011; Ram et al., 2018; Ram & Hadany 2012, 2014, 2019). In contrast, we also considered the possibility that the effects of the mutator loci are “self-referential”, that is, that the mutator loci govern their own mutation rate. This has a considerable impact on the ensuing evolutionary trajectories, as self-referential mutation rates can lead to a runaway effect: the higher the mutation rate, the faster it can change. Consequently, the evolution of higher constitutively expressed mutation rates (Figure 3) or of condition-dependent mutation rates (Figure 6) is sped up. However, this runaway effect also destabilises the evolutionary dynamics, leading to the “peaked” pattern seen in Figure 3A and Figure 6A, and increased variation between replicates. Why self-referential mutation rates are associated with improved adaptive tracking for constitutive mutation rates, but with worse adaptive tracking for condition-dependent mutation rates, is not entirely self-evident and may be the subject of future studies. However, we speculate that the constitutive mutation rate (encoded at a single locus) benefits from faster evolution. In contrast, the evolution of condition-dependent mutation rates involves more intricate coadaptation among three loci: the selection pressure on each locus depends on the current values of the other two loci. Thus, the condition-dependent mutation rate may be more strongly affected by the destabilising runaway effect observed under self-referential mutation rates, as this hampers the coadaptation of these three loci, leading to maladapted reaction norms and worse environmental tracking. Additionally, the plastic mutation strategy is encoded in three loci, and thus experiences a higher total mutation rate (i.e., the cumulative per-locus mutation rate of all three loci): this larger “mutational target” may make the plastic mutation strategy especially vulnerable, and more prone to mal-adaptation under high mutation pressure, which could contribute to the worsened adaptive tracking seen in the scenario of self-referential mutation rates.

The exploration of self-referential mutation rates is grounded in biology: mutation rates have been shown to vary strongly across the genome (Hodgkinson & Eyre-Walker, 2011; Monroe et al., 2022) – so genes that affect the mutation rate in one particular part of the genome without affecting the mutation rate of their own genetic basis likely exist (non-self-referential). However, some mechanisms that enhance the mutation rate, such as error-prone polymerases (Maslowska et al., 2019), will likely also affect the accuracy of replication of the genes in which they are encoded (self-referential). In nature, self-reference may also not be a categorical trait, as the extent to which the genome is affected by a mutation rate modifier likely varies continuously and depends on environmental conditions. However, since we show here that the evolutionary dynamics of self-referential and non-self-referential mutation rates differ in some respects, it is worth considering to what extent the mutation rate modifier considered in a particular study is self-referential or not - something that has been largely ignored in both the theoretical and empirical literature on mutation rates.

In general, our findings align well with those of Ram and Hadany (2012, 2019) and Lukačišinová and colleagues (2017), who, to our knowledge, report on the only other modelling approach concerning the evolution of condition-dependent mutation rates. Both studies conclude that condition-dependent mutation can spread or be maintained under a range of circumstances, as long as at least some mutations are beneficial (Ram & Hadany 2012) or there is sufficient need for repeated adaptation (Lukačišinová et al. 2017). Note that the modelling approach we used here is in many ways different: previous models have considered the spread of a genetic variant conferring a fixed condition dependence of the mutation rate in a population of individuals that express a constitutive mutation rate. Our model follows the evolution of a flexible condition-dependent mutation rate, allowing both condition-dependent and constitutive mutation strategies to emerge. Furthermore, in the model of Ram & Hadany (2012), the mutation rate responds to the presence of deleterious alleles, whereas in our model the mutation rate is linked to the mismatch of the ecological trait and the environment.

Despite this more flexible approach, our model makes several simplifying assumptions worth noting. For instance, we use only three evolving loci to model conditional strategies, which significantly restricts the shape of the function relating individual condition to the mutation rate. Furthermore, we focus on haploid organisms that reproduce asexually and exclude the effects of recombination. As previous studies have noted that recombination and sexual reproduction can disrupt mutation rate evolution by breaking the linkage between mutator loci and the mutations they induce (Martincorena & Luscombe, 2013; Sniegowski, et al. 2000; but see Cobben, et al. 2017 and Johnson 1999), the relevance of our results for sexually reproducing organisms thus still needs to be investigated. The high mutation rates that evolved in our models highlight another important simplification: an individual’s adaptation to its environment is determined by a single evolving locus. This facilitates a clearer analysis but also leads to an underestimation of the effect of deleterious mutations (e.g., we exclude traits under constant stabilising selection, which may experience a more adverse effect of elevated mutation rates). However, note that in nature, a phenotypic trait P is typically affected by many loci: the total mutation rate experienced by P may thus be quite high, even if the per-locus mutation rate is low.

Whether evolvability itself can evolve in response to selection for evolvability remains debated (Dawkins, 1988; Pigliucci, 2008; Payne & Wagner, 2019; Riederer et al., 2022; Pelabon et al., 2025). Previously, alternative explanations for the condition dependency of mutation rates have been proposed: individuals in poor condition may upregulate their mutation rate to increase evolvability, or to save energy by reducing the cost of replication fidelity. We here demonstrate that condition-dependent mutation rates not only enhance evolvability but also evolve in the absence of DNA replication costs, indicating that in our simulations the evolution of condition-dependent mutation rates is driven by selection for evolvability. We demonstrate that evolvability can itself be shaped by selection, leading to the evolution of condition-dependent mutation rates.

## Acknowledgements

We thank Ella Rees-Baylis, Luke Pattipeilohy and the members of the MARM group at the University of Groningen for discussions and input. We also thank the Center for Information Technology of the University of Groningen for their support and for providing access to the Perigrine/Hábrók high-performance computing clusters. F.J.W., T.J.B.v.E. and J.M.R. acknowledge funding from the European Research Council (ERC) under the European Union’s Horizon 2020 research and innovation programme (Grant agreement No. 789240); J.M.R. is supported by a GELIFES scholarship from the University of Groningen.

## Data and code availability

The C++ code for the simulations, the simulated data used to generate figures in this study, and the R code for data analysis can be found at https://doi.org/10.5281/zenodo.19663892.

