## Supplementary material for "The evolution of a condition-dependent mutation rate enhances evolvability"

### Contents

**Figure S1:** Evolution of condition-dependent mutation rates under intermediate selection strength.

**Figure S2:** Evolution of condition-dependent mutation rates under weak selection.

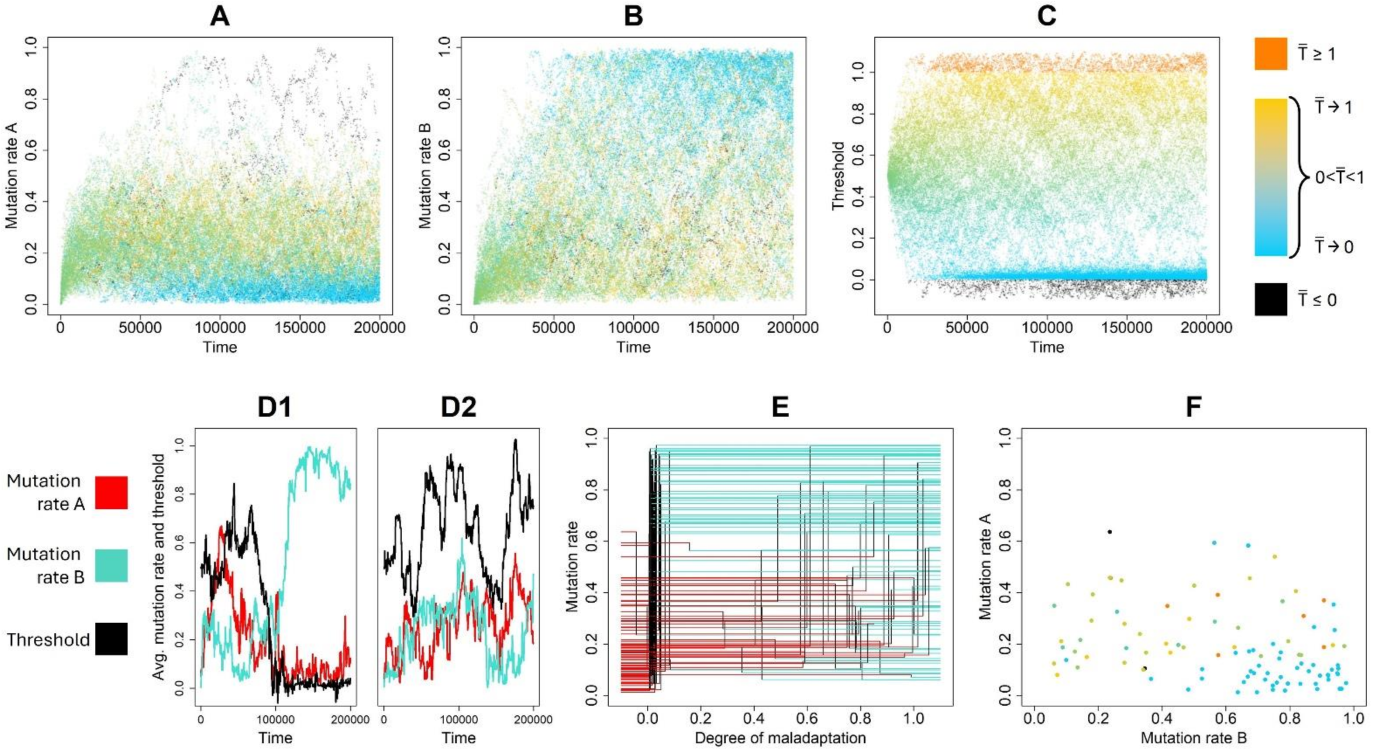


***Figure S1. Evolution of condition-dependent mutation rates under intermediate selection strength.*** *The figure corresponds in all aspects to Figure 4 in the main text, but now the selection strength is reduced from the default value s = 10 to s = 1, corresponding to a standard deviation of $\sqrt{0.5} = 0.707$ of the Gaussian fitness function. Average duration of environmental stasis D = 100. Under intermediate selection strength, condition-dependent mutation rates still emerge, but more slowly and (in the first 200,000 generations) not in all replicates. Panel D shows two representative examples: in one replicate population, a condition-dependent mutation rate similar to that observed in Figure 4 emerges after ca. 100,000 generations (D1), in the other replicate, it does not (D2).*

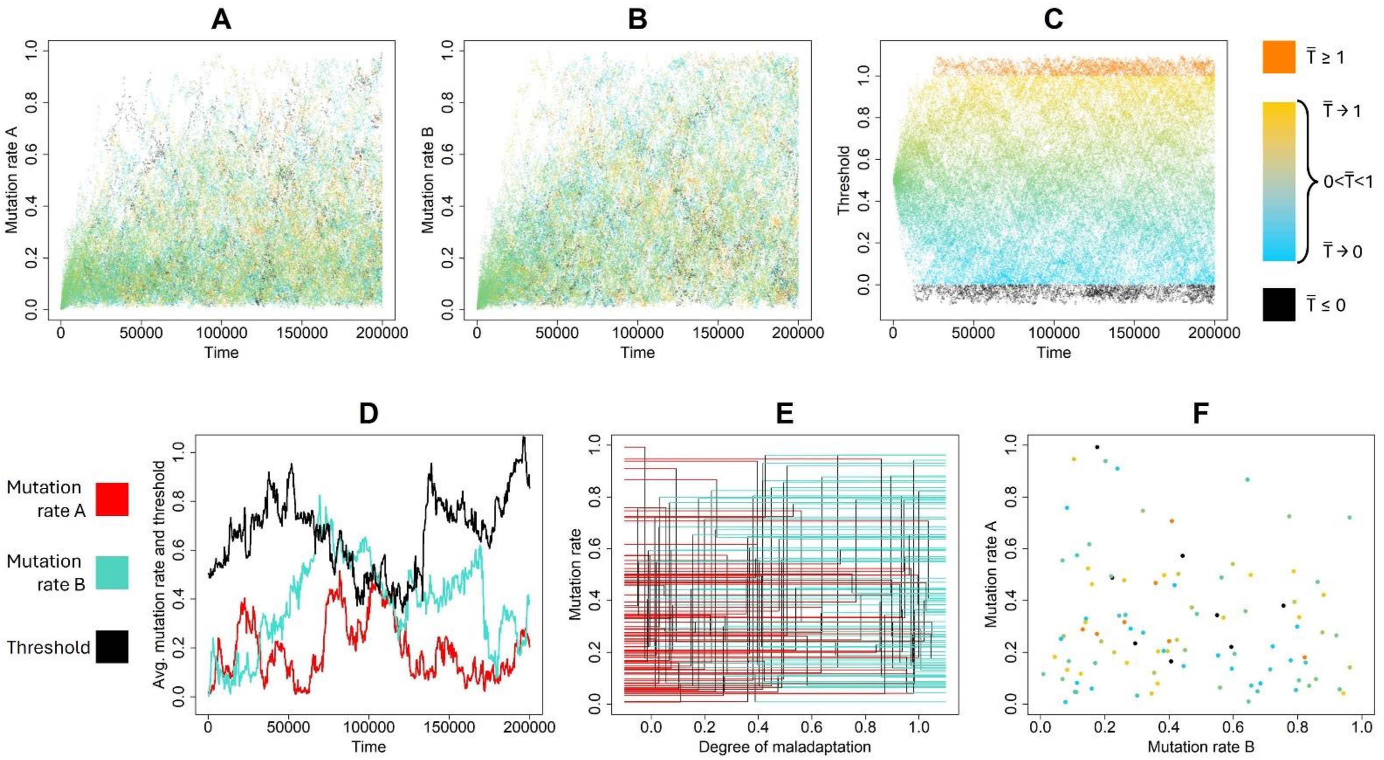


***Figure S2. Evolution of condition-dependent mutation rates under weak selection.*** *The figure corresponds in all aspects to Figure 4 in the main text, but now the selection strength is reduced from the default value s = 10 to s = 0.1, corresponding to a standard deviation of* $\sqrt{5} = 2.24$ *of the Gaussian fitness function. Average duration of environmental stasis D = 100. Under weak selection, a clear-cut condition-dependent mutation strategy (*$M_A$*,* $M_B$*, T) no longer evolves. Apparently, selection is too weak to overcome genetic drift.*
